# Significant inactivation of SARS-CoV-2 by a green tea catechin, a catechin-derivative and galloylated theaflavins *in vitro*

**DOI:** 10.1101/2020.12.04.412098

**Authors:** Eriko Ohgitani, Masaharu Shin-Ya, Masaki Ichitani, Makoto Kobayashi, Takanobu Takihara, Masaya Kawamoto, Hitoshi Kinugasa, Osam Mazda

**Affiliations:** Department of Immunology, Kyoto Prefectural University of Medicine, Kamigyo, Kyoto 602-8566, Japan; Central Research Institute, *ITO EN, Ltd.*, Makinohara, Shizuoka 421-0516, Japan

**Keywords:** Novel coronavirus, SARS-CoV-2, COVID-19, Tea catechin, Theaflavin, Polyphenols, Flavan-3-ols

## Abstract

Potential effects of teas and their constituents on SARS-CoV-2 infection were studied *in vitro*. Infectivity of SARS-CoV-2 was significantly reduced by a treatment with green tea, roasted green tea or oolong tea. Most remarkably, exposure to black tea for 1 min decreased virus titer to an undetectable level (less than 1/1,000 of untreated control). An addition of (-) epigallocatechin gallate (EGCG) significantly inactivated SARS-CoV-2, while theasinensin A (TSA) and galloylated theaflavins including theaflavin 3, 3’-di-gallate (TFDG) had more remarkable anti-viral activities. Virus treated with TSA at 500 μM or TFDG at 100 μM showed less than 1/10,000 infectivity compared with untreated virus. TSA and TFDG significantly inhibited interaction between recombinant ACE2 and RGD of S protein. These results strongly suggest that EGCG, and more remarkably TSA and galloylated theaflavins, inactivate the novel coronavirus.

## Introduction

The pandemic of novel coronavirus is expanding across the world with more than 65 million confirmed cases, and global death toll from COVID-19 has exceeded 1.5 million. Preventive and therapeutic procedures to conquer the virus infection is urgently needed. Repositioning of existing anti-viral agents and use of corticosteroid have markedly improved therapeutic outcome, while novel medicines including neutralizing monoclonal antibodies are being intensively developed.

We have explored food ingredients that inactivate SARS-CoV-2, because, if such materials are available, patients as well as healthy people can safely ingest them to eliminate virus from oral cavity and gastrointestinal tract and to prevent person-to-person transmission of the virus. Active components in such food ingredients may also be used as lead compounds to develop novel feasible and safe medicines to treat COVID-19 patients.

Teas, including green, black and oolong tea, reportedly have beneficial effects to human health, such as prevention of neoplasms as well as cardiovascular, metabolic, neurological, and infectious diseases ^1^ ^2^. Tea catechins are polyphenolic compounds contained in green tea and contribute to most of its biological effects ^3,4^. (-) epigallocatechin gallate (EGCG) is regarded a representative tea catechin with the highest activity. Tea catechins are quite unstable under certain environments such as alkalic solution and quickly form various multimers by oxidization. Theasinensin A (TSA) is a biologically active black and oolong tea polyphenol, produced by enzymatic oxidation of EGCG. It is one of the major compounds that are formed by incubating EGCG in culture medium ^5-7^. Theaflavins, typically theaflavin 3, 3’-di-gallate (TFDG), are contained in black tea at high concentrations, because catechins are converted into theaflavins by enzymatic oxidation during the manufacturing of tea leaves into black tea ^8 9^.

In the present study we examined whether SARS-CoV-2 can be inactivated by an exposure to tea. We also identified active components that contribute to the anti-viral effect.

## Results

To explore natural substances or food ingredients that inhibit SARS-CoV-2 infection, virus was treated with various food ingredient materials of botanical origins for 1 min, and the mixtures were subjected to TCID_50_ assays to determine viral titers (Supplementary Fig. S1A). As shown in Supplementary Fig. S1B, the viral titer fell to undetectable levels by the treatment with black tea, while Matcha green tea, roasted green tea (Hojicha) and oolong tea also decreased virus infectivity albeit to lesser extent.

In the following experiments, freeze-dried powders of each tea were used and osmotic pressure was adjusted by an addition of x2 DMEM. Virus was pretreated with each tea for 1 min, and the virus/tea/DMEM mixtures were immediately diluted 10-fold. After an addition of the diluted virus/tea/DMEM mixtures to the cells for 1 h, the supernatants were replaced by fresh culture medium (Fig. 1A). The TCID_50_ values indicated that black tea strongly inactivated the SARS-CoV-2, followed by green tea, Oolong tea and roasted green tea (Fig. 1B).

**Fig. 1.**
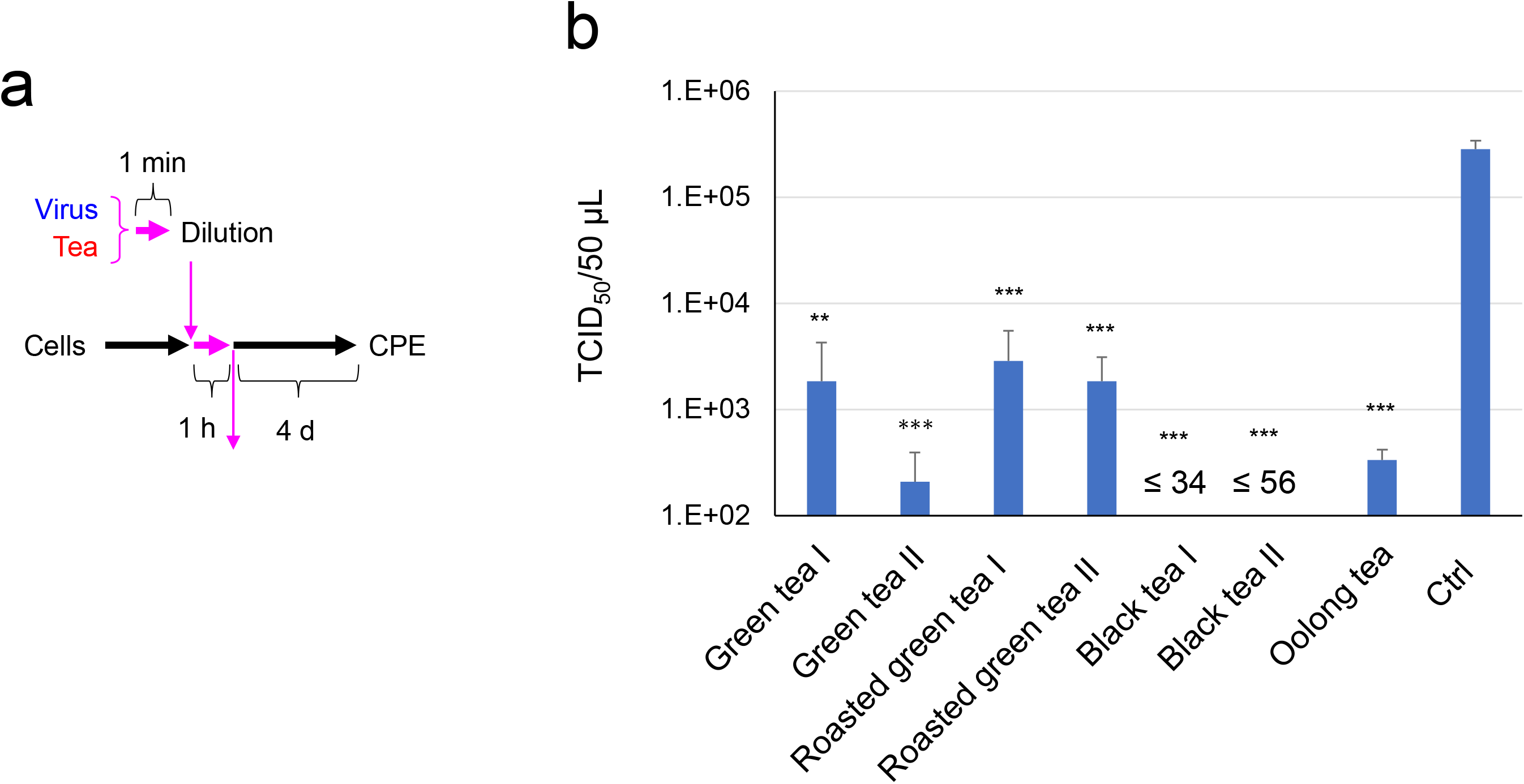
Infectivity of SARS-CoV-2 was significantly reduced by an exposure to tea. Virus was treated with ×1 concentration of the indicated tea/DMEM for 1 min, immediately followed by a 10-fold serial dilution and infection into VeroE6/TMPRSS2 cells to measure TCID_50_ values as described in the Materials and Methods. Scheme of experiments (a) and virus titer of each sample (b) are shown. Values are means ± S.D. (N=3). **P<0.01 and ***P<0.001 vs. Control.

Next, virus was treated with various doses of each tea, and after infection cell viability was estimated by WST-based analysis (Fig. 2A). Green tea, roasted green tea, black tea and oolong tea prevented SARS-CoV-2 from damaging VeroE6/TMPRSS2 cells in a dose-dependent fashion (Fig. 2B), confirming that teas significantly inactivated the virus.

**Fig. 2.**
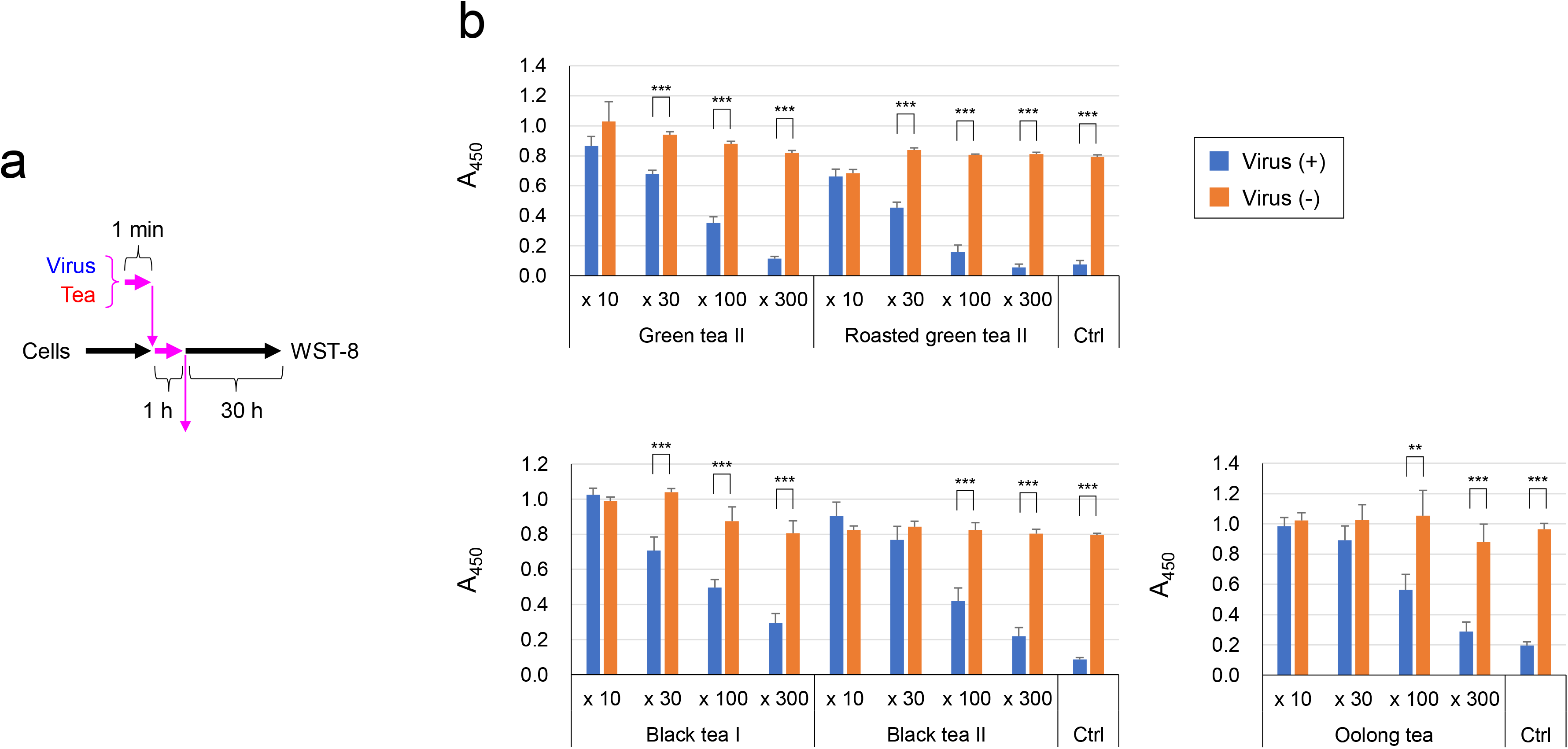
Viability of VeroE6/TMPRSS2 cells after infection with SARS-CoV-2 treated with tea. Virus was treated with each tea at the indicated dilutions for 1 min, and the virus/tea/DMEM mixture was added to VeroE6/TMPRSS2 cells at MOI=3 for 1 h. After removal of the supernatant, cells were cultured in fresh MS for 30 h, followed by cell viability determination as described in the Materials and Methods. Scheme of experiments (a) and A_450_ value of each sample (b) are shown. Values are means ± S.D. (N=3). **P<0.01 and ***P<0.001 between groups.

Then we tried to identify active components that are involved in the anti-virus function. We tested catechins and theaflavins that are well-known functional ingredients in teas (Supplementary Figs. S2 and Fig. 3A). As shown in Fig. 3B, TCID_50_ assay indicated that SARS-CoV-2 was significantly inactivated by EGCG but not by other catechins, i.e., (-) epigallocatechin (EC), (-) epicatechin gallate (ECG) and (-) epigallocatechin (EGC). We also prepared theasinensin A (TSA) and examined its effect on SARS-CoV-2. TSA more strongly inactivated the virus than EGCG did. Among theaflavins that we tested, theaflavin 3-O-gallate (TF3G), theaflavin 3’-O-gallate (TF3’G), and theaflavin 3,3’-di-gallate (TFDG) strongly inactivated the virus, whereas theaflavin (TF) failed to do so. The anti-virus effects of EGCG, TSA and TFDG were comparatively analyzed. As shown in Fig. 3C, TAAS and TFDG were more effective than EGCG in suppressing SARS-CoV-2, and the treatment with TSA at 500 μM or TFDG at 100 μM caused a significant drop in virus titer to undetectable levels (less than 1/10,000 compared with control).

**Fig. 3.**
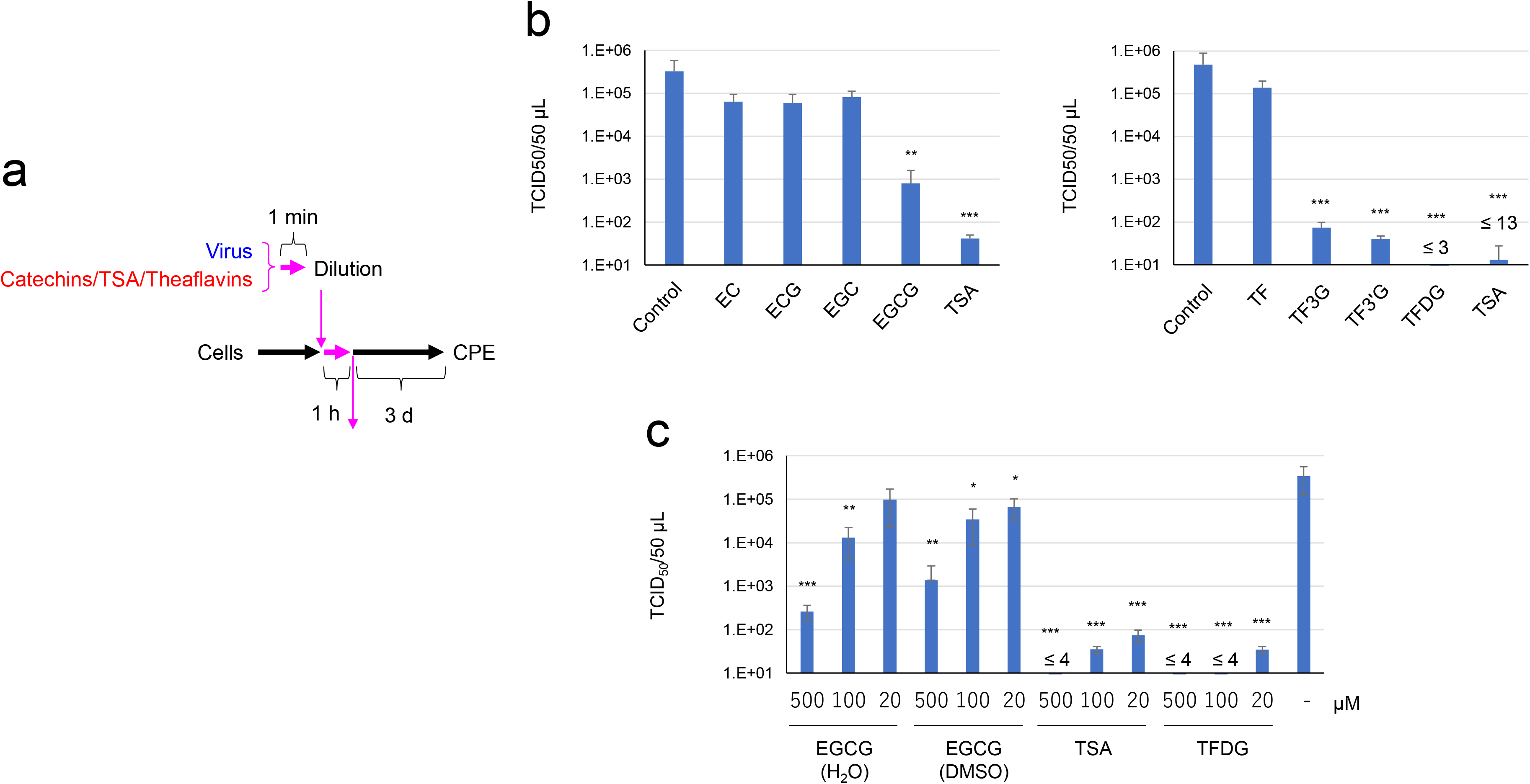
EGCG and galloylated theaflavins significantly decreased SARS-CoV-2 titer. Virus was treated with 500 μM (b) or the indicated concentrations (c) of tea catechin derivatives for 1 min, immediately followed by a 10-fold serial dilution and infection into VeroE6/TMPRSS2 cells to measure TCID_50_ values as in Fig. 1. Scheme of experiments (a) and virus titer of each sample (b and c) are shown. Values are means ± S.D. (N=3). *P<0.05, **P<0.01 and ***P<0.001 vs. Control.

These findings were confirmed by cell viability analysis, which clearly indicated that cell viability was not significantly reduced by infection with the TSA- or TFDG-treated virus. A higher concentration of EGCG was required to effectively protect cells compared with TSA and TFDG (Fig. 4A and B).

**Fig. 4.**
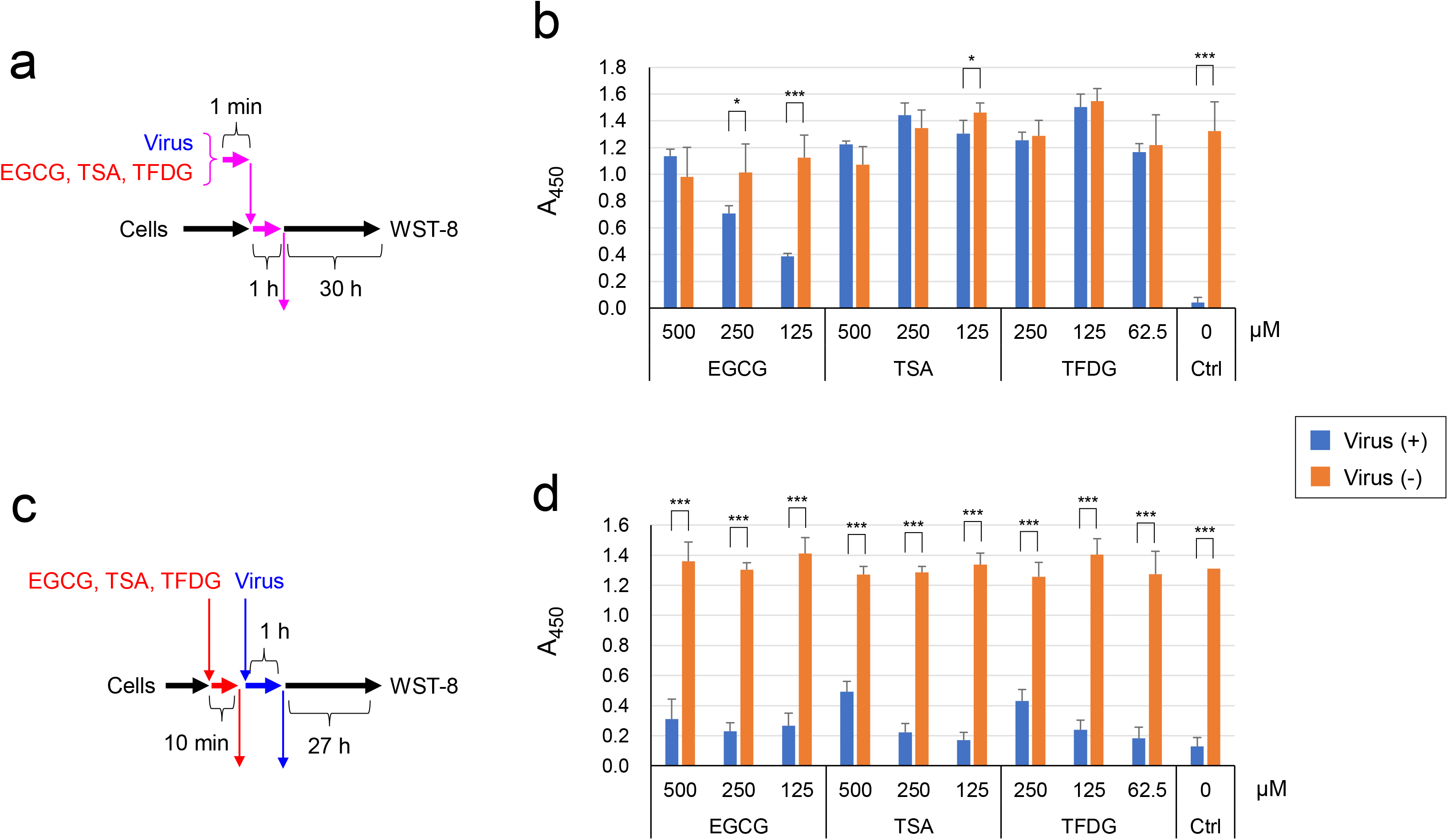
Catechin derivatives inactivated SARS-CoV-2, but didn’t render cells resistant to intact virus. a and b) Virus was treated with each tea catechin derivatives at the indicated concentrations for 1 min, and the mixture was added to VeroE6/TMPRSS2 cells at MOI=3 for 1 h. After removal of the supernatant, cells were cultured in fresh MS for 27 h, followed by cell viability evaluation as described in the Materials and Methods. c and d) Cells were pretreated with the indicated concentrations of the catechin derivatives for 10 min, followed by a removal of the supernatant, wash with PBS, and an addition of virus suspension (MOI=5). One h later, supernatant was replaced by fresh MS, and cells were culture for 27 h until cell viability was evaluated. Values are means ± S.D. (N=3). *P<0.05, and ***P<0.001 between groups.

In the next experiment, we preincubated cells with the three catechin derivatives for 10 min, followed by removal of the catechin-derivatives and infection with untreated virus (Fig. 4C). Cell viability analysis showed that the cells pretreated with EGCG, TSA, or TFDG were susceptible to virus as highly as non-treated cells were (Fig. 4D). This is in sharp contrast to the experiments using pretreated virus (Fig. 4B), strongly suggesting that the three catechin-derivatives may directly inactivate virions rather than interact with cell surface.

Real time RT-PCR analysis was performed to evaluate viral RNA in the culture supernatant and the cells that had been infected with the EGCG-, TSA- or TFDG-treated virus (Fig. 5A). As shown in Fig. 5B, treatment with 500 μM of TSA or TFDG resulted in significant suppression of RNA replication in the cells and virus release in the supernatant, while EGCG treatment did so less remarkably.

**Fig. 5.**
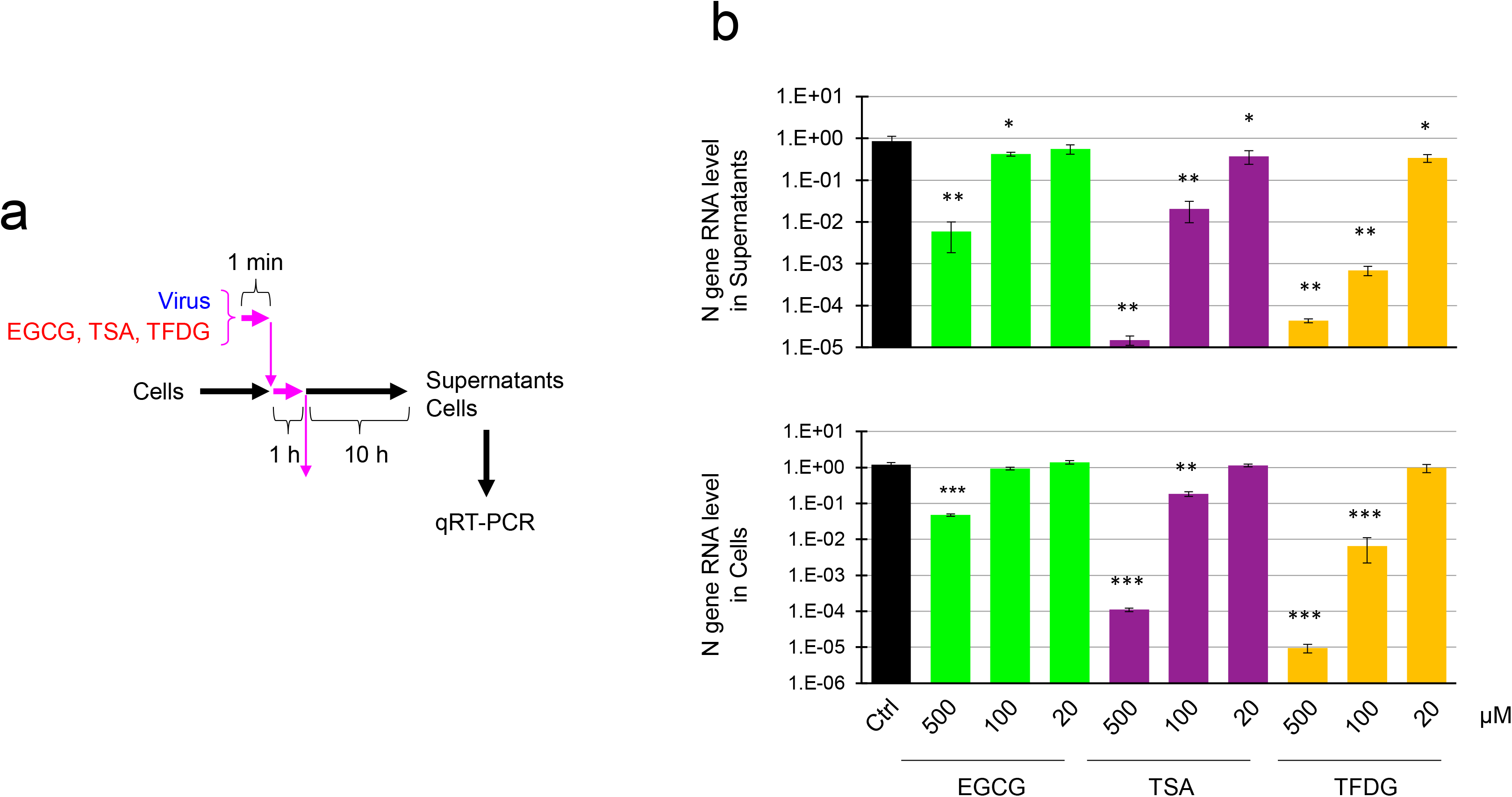
Evaluation of viral RNA in the supernatants and the cells infected with TSA- or TFDG-treated virus. Virus was treated with the indicated catechin derivatives for 1 min, and the mixture was added to VeroE6/TMPRSS2 cells at MOI=5 for 1 h. After removal of the supernatant, cells were washed with PBS and cultured in fresh MS for 10 h (a). RNA was extracted from the supernatants and cells, and real time-RT-PCR was performed as described in the Materials and Methods. Means ± S.D. of RNA levels are shown (N=4) (b). *P<0.05, **P<0.01 and ***P<0.001 vs. Control.

To figure out the mechanisms of inactivation of SARS-CoV-2 by EGCG, TSA and TFDG, we estimated influence of these catechin derivatives on the interaction between recombinant angiotensin converting enzyme 2 (ACE2) and receptor binding domain (RBD) of the SARS-CoV-2 spike protein. As shown in Fig. 6, TSA and TFDG strongly inhibited the interaction between these proteins, while EGCG also inhibited the protein interaction but to lesser extents than TSA and TFDG.

**Fig. 6.**
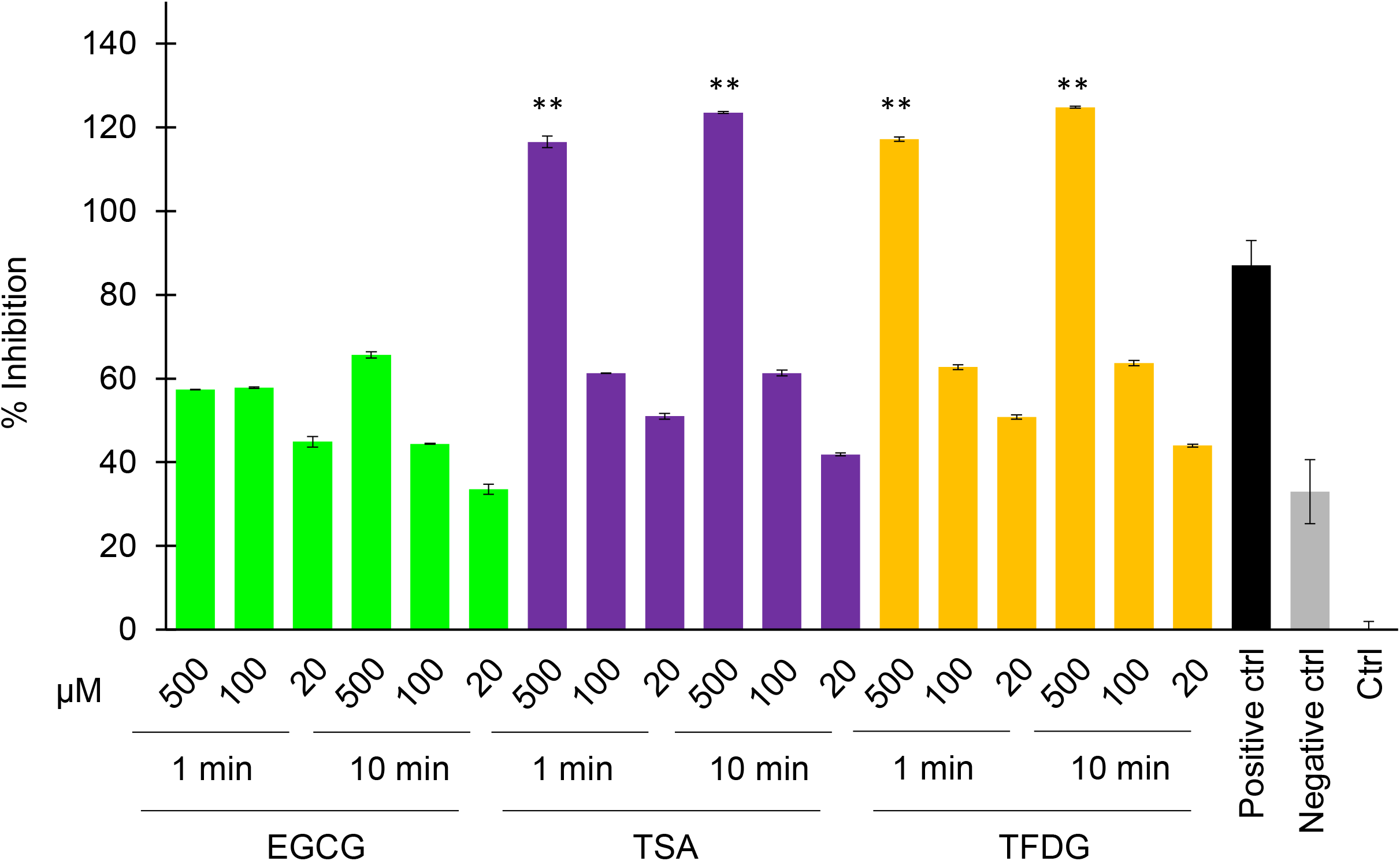
TSA and TFDG interfered with the interaction between ACE2 and RBD. Indicated catechin derivatives were subjected to a neutralizing assay using a SARS-CoV-2 Neutralizing Antibody Detection ELISA Kit (Cayman Chemical) as described in the Materials and Methods. %Inhibition for each sample is shown. Values are means ± S.D. (N=3 or 4). **P<0.01 vs. Negative control.

## Discussion

The present study demonstrated that EGCG inactivated SARS-CoV-2, while TSA and TFDG more significantly did so. Concentrations of tea catechins and theaflavins in tea beverages may considerably differ depending on the cultivars of tea plants, procedure and period of cultivation of the tea plants, levels of enzymatic oxidation during processing of tea leaves, etc ^10^. But green tea, oolong tea and black tea may contain enough high concentrations of EGCG, TSA and TFDG, respectively, for inactivation of SARS-CoV-2, considering the effective doses of the catechin derivatives shown in this study.

As for the mechanisms of the virus inactivation, our data strongly suggest that the catechin-derivatives may interact with virions rather than cells, because pretreatment of virion but not cells with catechin derivatives significantly suppressed infection (Fig. 4). Moreover, TSA and TFDG inhibited interaction between recombinant ACE2 and RBD (Fig. 6), strongly suggesting that these catechin derivatives may interact with RBD of S protein to prevent viral attachment to ACE2 on the cell surface. Our results are not consistent with previous in silico docking simulation studies that suggested high affinity binding of EGCG and non-galloylated theaflavin to RBD ^11,12^.

Tea catechins are reported to exert various biological effects including anti-oxidation, regulation of lipid metabolism, anti-inflammation, tumor suppression, and suppression of various viruses ^2-4^. Tea catechins are well known to suppress influenza virus infection by inhibiting attachment of the virions onto the cell surface ^4 13^. TSA has similar functions to EGCG, including antioxidant, anti-inflammatory, anti-obesity, anti-cancer and anti-bacterial functions ^14^. TFDGs are known to have anti-virus activity in addition to anti-obesity, anti-tumor, anti-atherosclerotic and anti-inflammatory functions ^15 16^.

So far as we know, etiological correlation between tea consumption and SARS-CoV-2 infection has not been reported. At the beginning of this study, we assumed that green tea may be most effective among the tea beverages, because green tea is ordinally and heavily consumed by a lot of people in Japan and Asian countries where remarkably less people have been infected with SARS-CoV-2 compared with Europe and North and Latin America ^10^. Unexpectedly, black tea that is consumed worldwide was more effective than green tea (Fig. 1). Thus, it remains unclear whether tea consumption influences on the incidence of SARS-CoV-2 infection.

SARS-CoV-2 may frequently undergo mutation, and it remains to be clarified whether the tea catechin-derivatives are effective in suppressing mutated virus.

Clinical significance and application of the present study should be carefully examined in future studies. After oral intake of tea, catechins and theaflavins are metabolized by intestinal microbiota and poorly absorbed and transported into blood stream in the gastrointestinal tract 17-20. Serum concentrations of catechins reach only quite low levels after a person ingest some cups of tea ^17,21^. The serum levels are substantially lower than the effective concentrations for suppression of SARS-CoV-2 infection (Figs. 3C and 4A). Nevertheless, oral intake of tea, EGCG, TSA and TFDG could inactivate SARS-CoV-2 in oral cavity and gastrointestinal tract. Infection at the gastrointestinal tract is regarded as an important characteristic of COVID-19 ^22–24^. Gargling with tea or catechin derivatives may also effectively block person-to-person transmission of the virus by eliminating virions from saliva. Tea is available at a large amount with low costs in broad area in the world including developing countries.

For many years, a lot of people ordinarily have tea beverages in the world, suggesting that teas and their constituents may be readily used as dietary-supplements, and potentially used as lead compounds to develop novel medicines with a relatively lower risk of causing severe toxic events compared with other chemicals. But it should be noted that intake of a large amount of catechins could cause adverse events including liver toxicity ^10 25^, while excess intake of caffein may also result in unbeneficial outcome.

## Materials and Methods

### Cells, virus, and culture medium

VeroE6/TMPRSS2 cells ^26^ were obtained from Japanese Collection of Research Biosources Cell Bank, National Institute of Biomedical Innovation (Osaka, Japan) and cultured in Dulbecco’s modified Eagle’s minimum essential medium (DMEM) (Nissui Pharmaceutical Co. Ltd., Tokyo, Japan) supplemented with G418 disulfate (1 mg/mL), penicillin (100 units/mL), streptomycin (100 μg/mL), 5% fetal bovine serum at 37°C in a 5% CO_2_ / 95% in a humidified atmosphere. The SARS-CoV-2 (Japan/AI/I-004/2020) were kindly provided from Japan National Institute of Infectious Diseases (Tokyo, Japan) and propagated using VeroE6/TMPRSS2 cells.

### Freeze-dried powders of tea extract

Tea extracts were prepared by soaking 40 g of ground up and homogenized tea leaves in 2,000 mL water at 80°C for 30 min. After centrifugation at 4,000 rpm for 15 min, supernatants were collected and filtrated through Toyo No. 2 filter papers, followed by evaporation and freeze-drying.

### Green tea catechins, theaflavins and catechin derivatives

EC, ECG, EGC, EGCG, TF, TF3G, TF3’G and TFDG were purchased from FUJIFILM Wako Pure Chemical Corporation (Osaka, Japan). TSA was synthesized from EGCG and purified as described. ^27^ Briefly, a solution of EGCG (1.0 g, 2.18 mmol) and CuCl_2_ (269 mg, 2.00 mmol) in 30% MeOH (400 mL) was vigorously mixed for 24 h. To the reaction mixture, ascorbic acid (10 g) was added and heated at 85 °C for 15 min. After chilling, the mixture was concentrated to evaporate MeOH and the resulting aqueous solution was applied to a Diaion HP20 column (Mitsubishi Chemical Corp., 3.0 cm i.d. ×25 cm) with water. After washing the column with water to remove reagents, sample was eluted with 70 % MeOH (300 mL), and applied to a preparative high-performance liquid chromatography (HPLC) using a YMC-Actus Triart C18 column (YMC CO., LTD., 20 mm i.d. × 250 mm). To elute the column, isocratic elution method was performed using a solvent mixture of Water:CH3CN = 3:2 at a flow rate of 16 mL/min. LC chromatograms were obtained at UV 280 nm. The fraction of TSA was collected and freeze-dried. The yield and purity of TSA were 251 mg and 96%, respectively.

### TCID_50_ assay for virus pretreated with tea

Freeze-dried powders of green tea, roasted green tea, black tea, and oolong tea was dissolved in sterilized distilled water at 78°C for 10 min to prepare x4 concentration of original tea. After chilling at room temperature, each solution was passed through a 0.45 μm filter, and placed into 96-well-plates at 50 μL/well in triplicate. Each sample was mixed with 50 μL of x2 serum-free DMEM, followed by an addition of 100 μL of SARS-CoV-2 suspension (1.5 × 10^6^ TCID_50_/50 μL) and incubation at room temperature for 1 min. Immediately, the virus/tea/DMEM mixture was serially diluted 10-fold with MS in the 96-well-plates. Chilled on ice, 100 μL of each sample was added to the VeroE6/TMPRSS2 cells that had been seeded into 96-well-plates at 5 × 10^4^/100 μL/well a day before. After culture for 4 days, cells were washed, fixed and stained with crystal violet solution to estimate CPE as described ^28^. TCID_50_ values were calculated by Reed-Muench method.

### TCID_50_ assay for virus pretreated with catechin derivatives

Solutions of catechins and theaflavins were diluted in MS (DMEM supplemented with 0.5% FBS) to concentrations of 2 mM, 400 μM, and 80 μM. The same volume of x2 DMEM was added to each solution followed by an addition of SARS-CoV-2 suspension (1.5 × 10^6^ TCID_50_/50 μL) and incubation at room temperature for 1 min. Immediately, the virus/catechin derivative/DMEM mixture was serially diluted 10-fold with MS, and TCID_50_ assay was performed as above.

### Cell viability assay for virus pretreated with tea

Tea was diluted and mixed with the same volume of serum free DMEM as above. Five μL of SARS-CoV-2 suspension (1.5 × 10^6^ TCID_50_/50 μL) or MS was added to 50 μL of each tea/DMEM mixture and incubated at room temperature for 1 min. Immediately, 50 μL of the virus/tea/DMEM mixture was added to the VeroE6/TMPRSS2 cells that had been seeded in 96-well-plates at a density of 5 × 10^4^/100 μL/well a day before (MOI=3 or 0). One hour later, culture supernatant was replaced by 100 μL of MS, followed by culture for 30 h. To measure cell viability, cells were washed with PBS, and WST-8 solution (Nacalai Tesque) diluted in phenol red-free DMEM was added to the wells at 50 μL/well. After culture for 45 min, 20 μL of 10% SDS was added to each well and OD at 450 nm was measured. Cell-free wells were regarded as a reference.

### Cell viability assay for virus pretreated with catechin derivatives

Catechin derivatives (EGCG, TSA and TFDG) was diluted and mixed with the same volume of x2 DMEM as above. SARS-CoV-2 suspension (1.5 × 10^5^ TCID_50_/50 μL) or MS was added to each mixture and incubated at room temperature for 1 min. Immediately, 50 μL of the virus/catechin derivatives/DMEM mixture was added to the VeroE6/TMPRSS2 cells that had been seeded into 96-well-plates at 5 x 10^4^/100 μL/well a day before (MOI=5 or 0). One hour later culture supernatant was replaced by fresh 100 μL MS, followed by culture for 27 h. Cell viability was measured as above.

### Cell viability assay using cells pretreated with catechin derivatives

Catechin derivatives/DMEM mixtures were prepared as above and added to the VeroE6/TMPRSS2 cells that had been seeded into 96-well-plates at 5 × 10^4^/100 μL/well a day before. Ten min later, supernatants were discarded, and wells were washed with PBS, followed by an addition of SARS-CoV-2 suspension (1.5 × 10^5^ TCID_50_/50 μL) to the wells for 1 hour (MOI=5 or 0). After replacement of the culture supernatant by 100 μL of MS, cells were cultured for 27 hours, and cell viability was measured as above.

### Real time-RT-PCR

RNA was extracted from culture supernatant using TRI Reagent® LS (Molecular Research Center, Inc., Montgomery Road, Cincinnati, OH, USA) and reverse-transcribed using ReverTra Ace^®^ qPCR RT Master Mix (Toyobo, Shiga, Japan). Quantitative real-time PCR was performed using a Step-One Plus Real-Time PCR system (Applied Biosystems, Foster City, CA, USA) and the following set of primers/probes specific for viral N gene. Forward primer, 5’-AAATTTTGGGGACCAGGAAC-3’; reverse primer, 5’-TGG-CAGCTGTGTAGGTCAAC-3’; and probe, 5’-(FAM) ATGTCGCGCATTGGCATGGA (BHQ)-3’.

### Neutralizing assay

Neutralizing assay was performed using SARS-CoV-2 Neutralizing Antibody Detection ELISA Kit (Cayman Chemical, Ann Arbor, MI, USA; Item No. 502070) according to the manufacturer’s protocol.

### Statistical analysis

Statistical significance was analyzed by Student’s *t* test, and P<0.05 was considered significant.

