## Supplementary Figs. 1-2 for "Significant inactivation of SARS-CoV-2 by a green tea catechin, a catechin-derivative and galloylated theaflavins *in vitro*"

### Supplementary Fig. S1

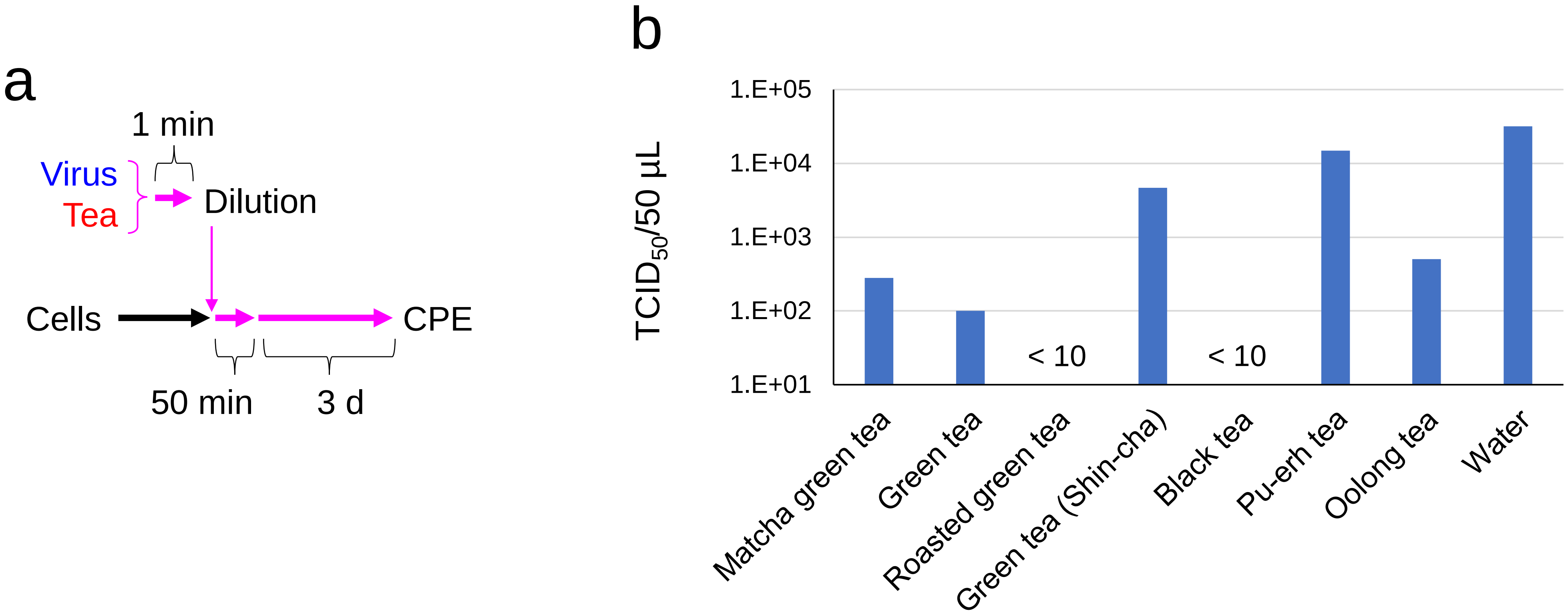


Supplementary Fig. S1

Powdered *Matcha* green tea, as well as leaves of green tea, roasted green tea (*Hojicha*), green tea (*Shin-Cha*, i.e., first picking of a season), black tea, Pu-erh tea and oolong tea were purchased at a supermarket in Kyoto, and brewed in hot water according to each recipe indicated on the package. After filtrated through a 0.22 μm filter, 500 μL of each sample was mixed with 20 μL of SARS-CoV-2 suspension (5 x $10^5$ $TCID_{50}$/50 μL) and incubated at room temperature for 1 min. Immediately, each mixture was serially diluted 10-folds into MS, and 50 μL of the samples were added to VeroE6/TMPRSS2 cells that had been cultured in 96-well-plates overnight (N=4 wells per each dilution). Plates were incubated for 50 min with gentle shaking every 10 min. Fifty μL of MS was added to the cells, which were subsequently cultured under standard conditions for 3 days. Cells were stained with Crystal Violet, and $TCID_{50}$ values were calculated by Reed-Muench method as described elsewhere (N=4 wells per each dilution). a) Experimental protocol is shown. Pink arrows represent the presence of tea and virus. b) $TCID_{50}$/50 μL values are shown.

### Supplementary Fig. S 2

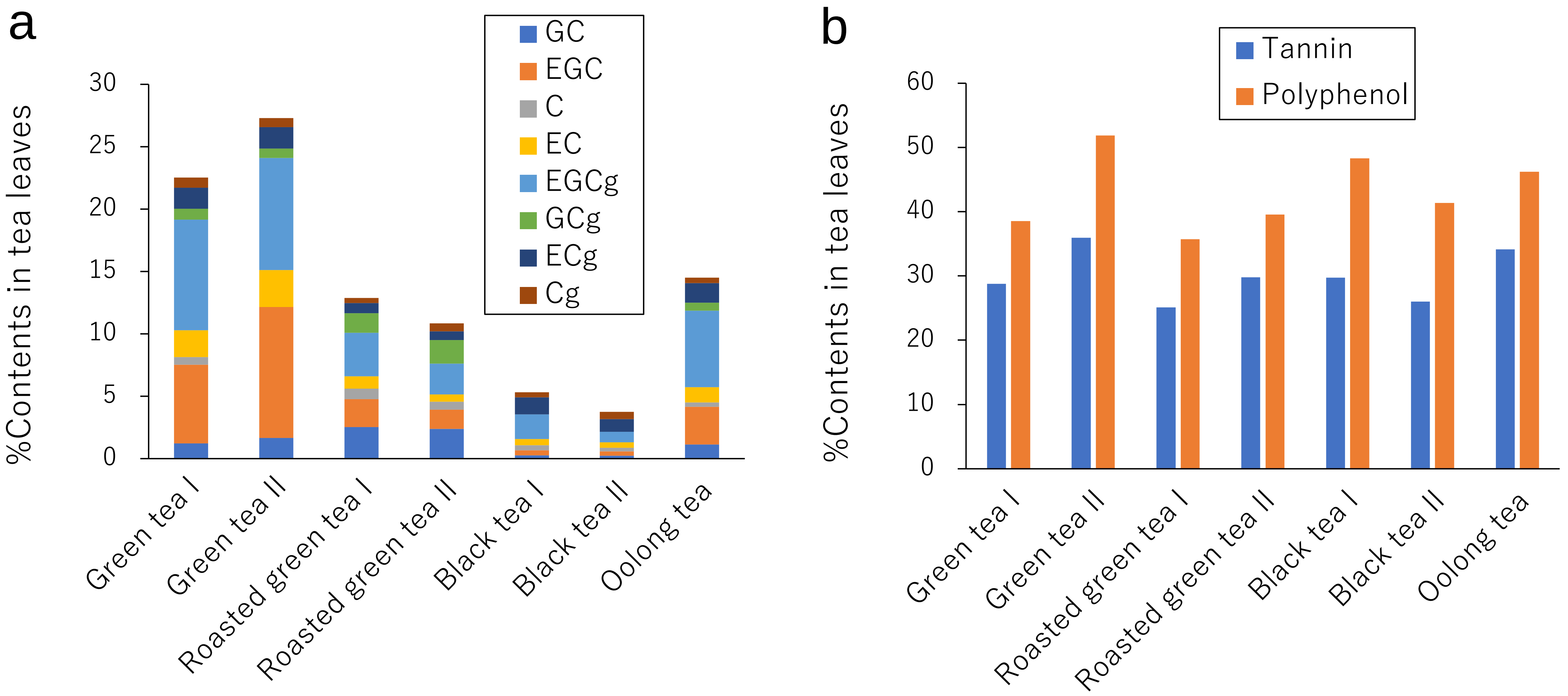


Supplementary Fig. S2
(a) %Contents of catechins (b) as well as tannin and polyphenol (B) in each tea were determined.
